# Human rhinovirus 16 infection modifies the microRNA landscape during epithelial cell infection

**DOI:** 10.1101/2023.12.08.570683

**Authors:** Pax Bosner, Victoria Cappleman, Alka Tomicic, Aref Kyyaly, Jamil Jubrail

**Author notes:** Correspondence: Jamil Jubrail.

## Abstract

Human rhinovirus is a major driver of disease exacerbations for patients with asthma and chronic obstructive pulmonary disease. Importantly, the virus can disrupt immune cell functioning leading to secondary bacterial infections in these patients. MicroRNAs are important regulators of gene expression and control viral pathogenesis and immune cell functions. However, the role of different miRNAs during rhinovirus infections remains poorly defined. The aim of this study was to analyze the impact of RV16 on epithelial cell responses and identify miRNAs with altered expression levels in response to the viral infection and their target genes. HeLa-Ohio cells exposed to RV16 showed deficiencies in bacterial clearance and induced changes in the expression of several novel miRNAs identified using a custom panel of 88 different miRNAs and further analyzed using our newly developed computer program. Our experiments identified a panel of 9 differentially regulated miRNAs and our computer program narrowed this down to two that could represent biomarkers for RV16 infection. Our results also showed that upon upregulation of these miRNAs (miR-101-3p and miR-30b-5p) by RV16 there was a downregulation in EZH2, RARG and PTPN13 that could correlate with a productive viral infection and a worsened immune response. Taken together, our findings suggest that RV16 modifies the miRNA landscape in epithelial cells and that miR-101-3p and miR-30b-5p could represent key biomarkers for either early or late RV16 infection.

## Introduction

Rhinovirus is a member of the *Picornaviridae* family and is a positive sense, single stranded RNA virus of approximately 7,200bp and has a complex life cycle in multiple cell types [1-3]. This complex lifecycle allows the virus to infect different cells within the respiratory tract with different efficiencies and cause a wide spectrum of respiratory tract infections. Although predominantly thought of as an upper respiratory tract infection (URTI), RV can cause severe lower respiratory tract disease in patients with inflammatory airway diseases such as asthma [4-5]. Numerous studies have implicated RV as a primary causative pathogen causing severe lower airway disease during exacerbations that can then lead to the development of secondary bacterial infections [6]. The underlying mechanisms behind this remain to be fully resolved with only a handful of mechanistic studies published [7-9]. Within these studies individual protein players were identified as being modified upon RV infection but whether they correlate with RV infection has never been fully determined. To address this, we sought to establish a global screen of RV infected epithelial cells to identify potential miRNA players that could link innate immune changes with ongoing RV infection.

miRNAs are small non-coding RNAs that are roughly 19 to 25 nucleotides long and regulate gene expression transcriptionally and post-transcriptionally and therefore integrate and control numerous cellular processes [10]. Within the last two decades, miRNAs have gained rapid interest owing to their potential to be biomarkers of multiple diseases. Our group recently demonstrated their importance in asthma pathogenesis which is exacerbated by RV [10-12]. In recent years, whether miRNAs could be markers of viral infections has been examined with a lot of correlative data, but no clear pattern observed [13-14]. In RV infections numerous studies have implicated miRNAs as potential biomarkers but these have been done with limited panels, at one time point or in patients with underlying diseases [15-16]. This makes the validity of the findings difficult to interpret although they do clearly demonstrate the importance of miRNA changes upon RV infection.

In this study we sought to determine whether specific miRNAs were modified in epithelial cells upon early and late RV infection. Our results clearly found that there was a differential pattern of miRNA expression with differences at different time points of infection. From these results we can suggest that a subset of miRNAs could represent markers to screen at risk individuals with a further subset useful as potential diagnostic biomarkers for ongoing infection.

## Materials and Methods

### Cell culture

HeLa-Ohio cells were sourced from the European Collection of Authenticated Cell Cultures (ECACC). HeLa-Ohio cells were grown in DMEM supplemented with 10% foetal-calf serum, 1% L-glutamine and 1% penicillin-streptomycin. Cells were maintained and passaged as previously described [8]. HeLa-Ohio cells were used in this study because they demonstrate a good propensity for infection by numerous rhinovirus strains and can be easily manipulated to allow infection of rarer serotypes. Most studies of rhinovirus use either HeLa-Ohio cells or HeLa-H1 cells initially and work performed in these cells is often reproduced in various cell lines and primary cells as we have found in the past [8, Jubrail *et al*., 2023, in revision].

### Human rhinovirus production and quantification

Human rhinovirus 16 (RV16) VR283 was purchased from the American Type Culture Collection (ATCC) and produced from HeLa-Ohio cell lysates as previously described [8]. Briefly, HeLa-Ohio cells at a density of 1 × 10^6^ were infected with 1 ml pure RV16 in 2 ml virus medium (DMEM supplemented with 4% foetal-calf serum and 1% L-glutamine). Cultures were placed at room temperature on a 50-rpm shaker for 1 hour and then inoculated with infection medium and left at 37°C, 5% CO2 for 48 hours. Virus was then produced by freeze-thawing, centrifugation and filtration and stored in aliquots at - 80°C as previously outlined [8]. All stocks were quantified using a tissue culture infective dose 50 (TCID50) assay and plaque assay as previously outlined [17-18].

### Growth of bacteria

*Escherichia coli* or *Staphylococcus aureus* were grown in 10 ml LB agar overnight by taking 1 colony from a streaked-out plate. The next day the OD600 was measured, and the bacteria were diluted in phagocytosis medium (DMEM with 1% L-glutamine) to the correct multiplicity of infection (MOI) and used in downstream experiments.

### Assessment of bacterial internalization

HeLa-Ohio cells were challenged with RV16, ultraviolet (UV)-inactivated RV16 or mock medium at an MOI 1 for 1 hour at room temperature followed by an overnight rest. The next day cultures were washed with phosphate buffered saline (PBS) and then challenged with either *E. coli* or *S. aureus* at an MOI 5 for 1 or 2 hours respectively. At each time point cultures were washed with PBS and treated with 20 μg/ml gentamicin for 30 minutes before being washed with PBS and incubated with 2% saponin for 12 minutes. Cultures were then lysed vigorously and plated onto LB agar to enumerate intracellular bacterial colony forming units.

### Assessment of bacterial clearance

HeLa-Ohio cells were challenged with RV16 or mock medium as above and the next day incubated with *S. aureus* as above. After 2 hours, cultures were treated in the same way as above and after 30 minutes either washed and incubated with 2% saponin or washed and maintained in 2 μg/ml gentamicin for 30 minutes, 2 hours or 4 hours. At each of these time points cultures were lysed with saponin as above and plated onto LB agar to enumerate intracellular bacterial colony forming units.

### MicroRNA purification and extraction

MicroRNA was purified and extracted using the Qiagen miRNEASY Tissue/ Cells Advanced Kit. Briefly, the samples were first thawed at 37°C for 10 minutes and then centrifuged at 14000 x g for 15 minutes. 200 μl was then transferred into a 2 mL microcentrifuge tube, to which 60 μL of buffer RPL was added. The samples were vortexed at 44 *x* g for 5 seconds each. The samples were then incubated at room temperature for 3 minutes. After incubation,1 μL of spike-in mix was added followed by 20 μL of buffer RPP and samples were vortexed again at 44 *x* g for 20 seconds. All samples were then left at room temperature for 3 minutes and were then centrifuged at 12000 x g for 3 minutes. The supernatants were transferred into new microcentrifuge tubes and 220 μL of isopropanol was added. The tubes were vortexed for 10 seconds at 100 x g. The samples were then transferred into a RNeasy UCP MinElute column and centrifuged at 8000 x g for 15 seconds. Then 700 μL of RWT was added into the spin column and they were centrifuged for 15 seconds at 8000 x g. 500 μL of 80% ethanol was added to the tubes and they were centrifuged for 2 minutes at 8000 x g. The columns were placed into collection tubes and centrifuged at 13586 x g for 5 minutes, with the lids open. The columns were then placed into new collection tubes and 20 μL of RNase-free water was directly added to the centre of the spin columns and they were then incubated at room temperature for 1 minute. The columns were placed back into the centrifuge for 1 minute at 13586 x g. The collection tubes, with the total RNA were placed into the freezer at -80°C until further use.

### Reverse transcription and qPCR

The reverse transcription (RT) reaction mixture consisted of 2 μL of miRCURY SYBR Green reaction buffer, 4.5 μL of RNase-free water, 1 μL of 10x miRCURY RT Enzyme Mix, 0.5 μL of UniSp6 RNA spike-in and 2 μL of the template RNA. For complete fusion of the product, the samples were briefly vortexed for 10 seconds The reaction was then run through a program of 40 cycles according to the program in Table 1:

**Table 1.** Conditions used for reverse transcription.

| Setting | Temperature (°C) | Duration (mins) |
| --- | --- | --- |
| Priming | 42 | 60 |
| Denaturation | 95 | 5 |
| Storage | 5 | Up to 4 days in fridge |

Human Inflammatory Response & Autoimmunity Focus) (Qiagen, YAHS-205Z) were used for each of the different samples. The reaction mixture for the qPCR reaction included 500 μL of 2x miRCURY SYBR Green Master Mix, 5 μL of undiluted cDNA template and 495 μL of RNase-free water. Each reaction mixture was briefly vortexed for 10 seconds before distributing 10 μL of the mixture across the entire plate. The plate was sealed and left at room temperature for 5 minutes. The qPCR program ran through 40 cycles according to the real-time cycler setup in table 2:

**Table 2.** qPCR reaction template.

| Setting | Temperature (°C) | Duration (seconds & mins) |
| --- | --- | --- |
| Initial Heat Activation | 95 | 120 minutes |
| Denaturation | 95 | 10 seconds |
| Annealing and Extension | 56 | 60 seconds |

The data produced from the real-time cyclers for each sample was stored on the Qiagen Software 96, where all files were uploaded onto.

### MicroRNA analysis

MicroRNA analysis was performed using GeneGlobe. GeneGlobe is an online portal that takes the qPCR data and uses an algorithm to normalise the data, run quality control analysis, and identify the up and downregulated miRNAs and confirm the samples are suitable for analysis. In our experiments we used GeneGlobe to normalise the data and perform statistical analysis to determine the significance in the changes of the expression of the miRNAs and correlate this to expression levels all compared to internal controls. This produced a detailed report that outlined all the different up and downregulated miRNAs.

### Development of program

The developed program was coded using Python and the chosen IDE was Spyder – available at https://www.spyder-ide.org/. The specific version of IDE used in this project was Spyder version 5, running on Python version 3.9. Hardware used for program development and running was Raspberry Pi 4 (model B) computer; CPU - Broadcom BCM2711, Quad core Cortex-A72 (ARM v8) 64-bit SoC @ 1.8GHz; RAM - 4GB LPDDR4-3200 SDRAM. The program interacts with two databases, mirDIP and VarElect, through API requests. MirDIP is a miRNA Data Integration Portal, which links miRNAs with their target prediction genes. The current database version 5.2.3.1 was used in this project [19]. VarElect is a bioinformatics tool that can identify direct and indirect links between an array of genes and a set phenotype, and rank the genes based on the strength of that relationship [20]. The program development started with the importing of the modules required for formatting and interacting with the databases. Using the class *mirDIP_Http*, the calling point for the mirDIP database and the parameters query, as well as the input parameters (miRNAs), an API request was sent. The mirDIP database returned the genes and scores of the miRNA-gene relationship. Those results were extracted and formatted into a DataFrame and exported as a csv file. Furthermore, the program used the data (list of genes) received from the mirDIP database to set up the parameters, alongside the phenotype of interest, and sent an API request to the VarElect database. The VarElect database returned the ranked genes that have a direct relationship to the set phenotype, together with p-value and a gene-phenotype relationship score. Those results were also extracted and formatted into a DataFrame and exported as a csv file.

### Statistics

All statistics were performed using GraphPad Prism version 9 and are stated in the figure legends. Statistical testing was done via a non-parametric Two Way Anova. Statistical significance was determined using appropriate post-tests versus control conditions and significance accepted when p < 0.05. All post-tests are stated in the figure legends.

## Results

### HeLa-Ohio cells robustly produce RV16

We first wanted to determine whether HeLa-Ohio cells could produce infectious RV16 24 hours post-infection. We challenged HeLa-Ohio cells with different MOIs of RV16 or mock medium for 1 hour at room temperature followed by 23 hours rest. The next day supernatants were collected and titrated onto fresh HeLa-Ohio cells and viral production measured as both the number of wells with at least 50% dead cells (figure 1A-B) and by determining the PFU/ml (figure 1C-D). We found that at all MOI HeLa-Ohio cells were producing RV16 that was above the input viral inoculum (figure 1). Our results showed that this was dose dependent with MOI 0.1 showing the least virus production with ∼99% of wells still showing greater than 50% of alive cells and a viral titre of ∼ 5 × 10^7^ PFU/ml (figure 1) and MOI 10 showing the greatest virus production with ∼25% of wells still showing greater than 50% of alive cells and a viral titre of ∼ 2 × 10^9^ PFU/ml (figure 1). Interestingly, we found MOI 0.2-1 showing similar numbers of wells with greater than 50% alive cells (figure 1A-B) and MOI 0.2-3 showing a similar PFU/ml (figure 1C-D). We also used these supernatants in downstream tests and showed the virus produced was infectious (data not shown). Taken together, these results demonstrate that when challenged with RV16, HeLa-Ohio cells can sustain viral infection and produce infectious viruses.

**Figure 1:**
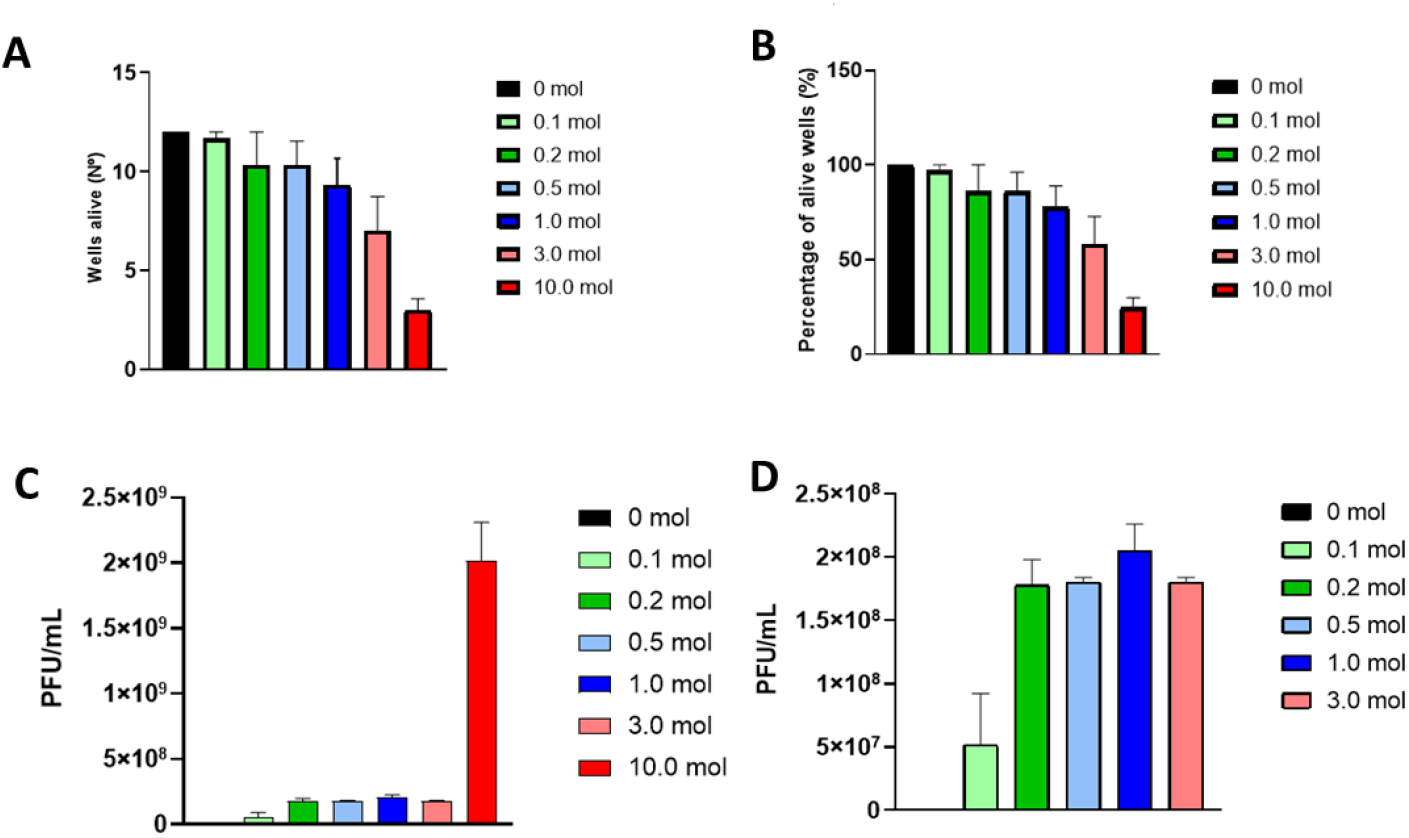
HeLa-Ohio cells produce RV16 after 24 hours. HeLa-Ohio cells were challenged with RV16 or mock medium for 24 hours and then supernatants collected and viral titres determined. (A) Number of wells with greater than 50% cells alive at each MOI, (B) Percentage of wells with greater than 50% cells alive at each MOI, (C) PFU/ml at each MOI, (D) PFU/ml with MOI 10 excluded. n=3, error bars represent SEM.

### RV16 infection does not impact bacterial phagocytosis in HeLa-Ohio cells

We next wanted to determine if HeLa-Ohio cells infected with RV16 would show an impairment in their ability to internalise bacteria. We challenged HeLa-Ohio cells with RV16 at an MOI 1 as above and after 24 hours exposed them to either E. coli or S. aureus at an MOI 5. We measured bacterial internalisation after 1 hour (E. coli) or 2 hours (S. aureus) following washing and treatment of cultures with antibiotics to kill non-internalised extracellular bacteria. We found that compared to our previous work with macrophages [8], RV16 infection of HeLa-Ohio cells did not cause any significant impairment in bacterial internalisation (Figure 2). This suggests that the internalisation machinery in HeLa-Ohio cells following RV16 infection is not perturbed.

**Figure 2:**
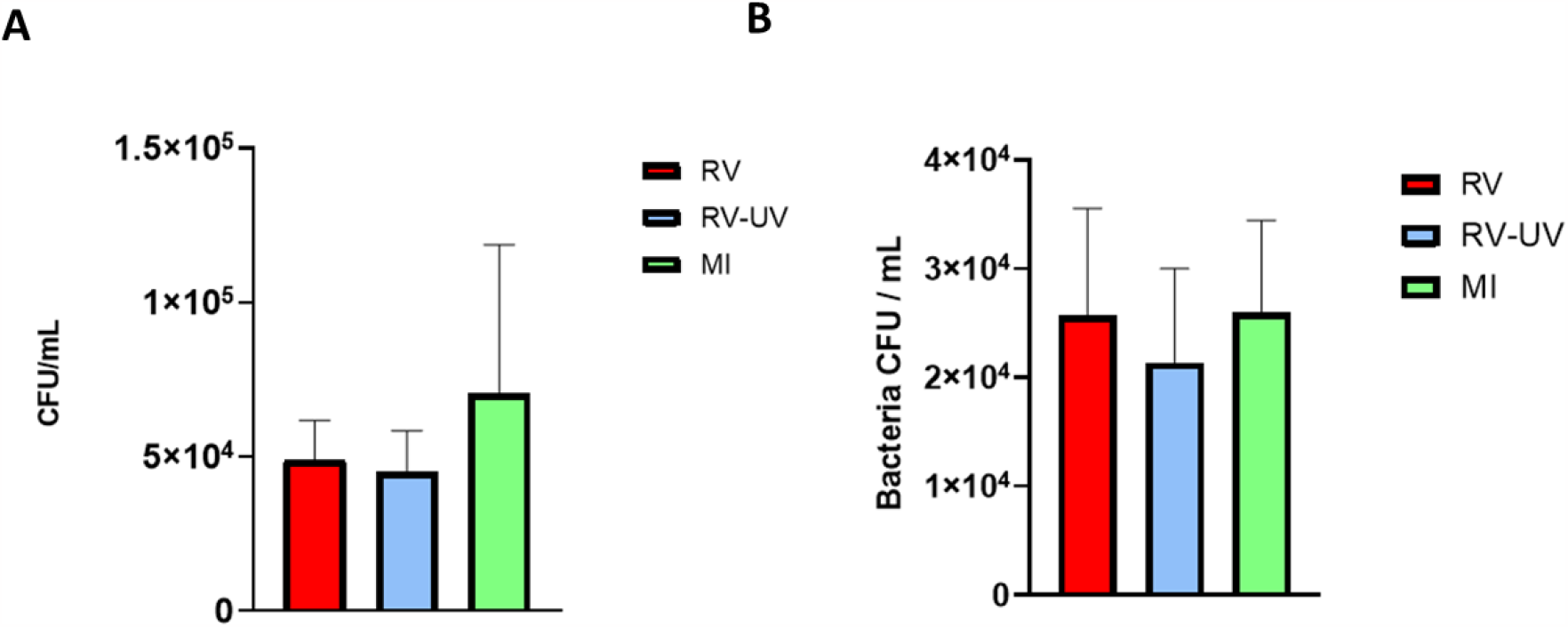
RV16 does not impact bacterial internalisation in HeLa-Ohio cells. HeLa-Ohio cells were challenged with RV16, UV-inactivated RV16 or mock medium at an MOI 1 for 24 hours and then exposed to bacteria for up to 2 hours. (A) *E. coli* internalisation and (B) *S. aureus* internalisation. n=3, error bars represent SEM.

### RV16 does not impact bacterial clearance

Following on from the above we next wanted to determine whether RV16 impacted HeLa-Ohio cells ability to clear intracellular bacteria. We challenged HeLa-Ohio cells with RV16 at an MOI 1 as above and after 24 hours exposed them to S. aureus at an MOI 5 for 2 hours. Following removal of non-internalised extracellular bacteria, intracellular bacterial numbers were followed over 4 hours by maintaining cultures in a low dose of antibiotic and lysing cells at each time point. Our results showed that by looking at the CFU/ml alone there appeared to be no significant difference in intracellular bacterial clearance between RV16 infected or mock infected conditions (Figure 3A). When we converted these CFU/ml values into a percentage we found that the infected cultures showed a faster percentage decrease in intracellular CFU/ml compared to mock cultures within the first 30 minutes which was not sustained (figure 3B). By 4 hours whilst mock infected cultures had nearly eliminated the intracellular bacteria, we found that infected cultures showed an overall increase in bacterial numbers (figure 3). These results suggest that while RV16 infected HeLa-Ohio cells can clear intracellular bacteria initially, they exhaust this ability over time creating a niche for intracellular staphylococcal replication.

**Figure 3:**
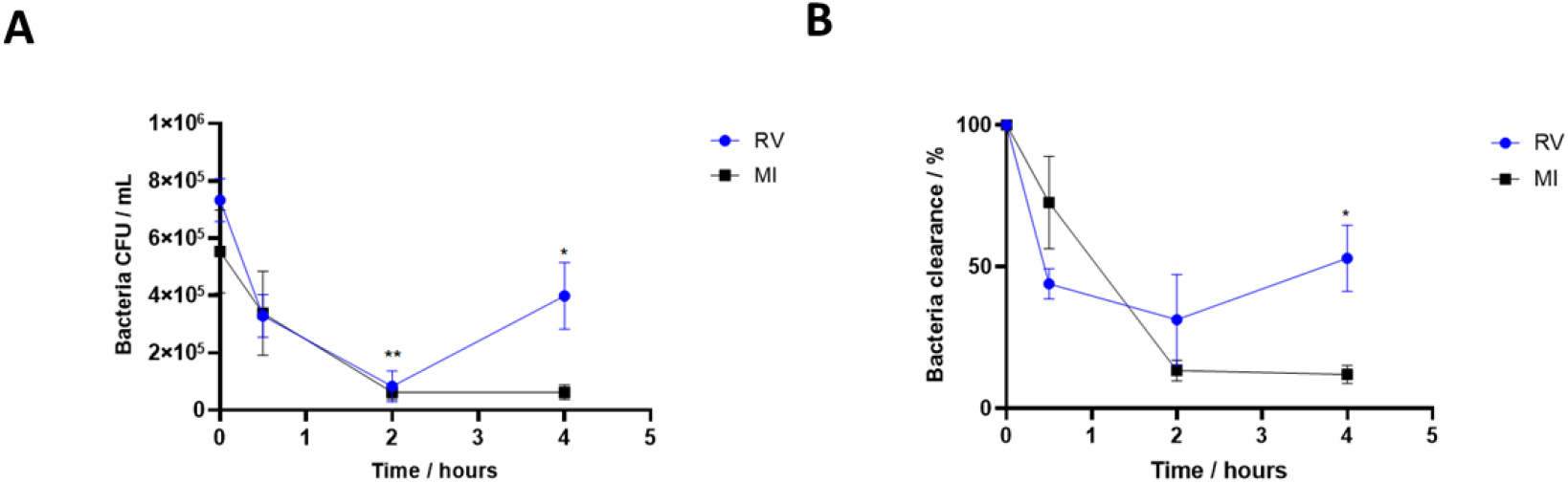
RV16 infected HeLa-Ohio cells cannot clear intracellular *S. aureus*. HeLa-Ohio cells were challenged with RV16, or mock medium at an MOI 1 for 24 hours and then exposed to *S. aureus* for 2 hours and intracellular bacterial clearance followed over 4 hours. (A) CFU/ml over time and (B) Percentage change in intracellular bacterial numbers. n=3, error bars represent SEM. *p<0.05, **p<0.01, Two Way Anova with Dunnett’s post test vs time 0.

### RV16 infection alters the micro RNA landscape of HeLa-Ohio cells

Based on the above findings we next wanted to determine if there were any global changes in RV16 infected epithelial cells that could identify potential mechanisms behind the inability of HeLa-Ohio cells to eradicate intracellular S. aureus. To do this we decided to analyse miRNA expression which could allow us to identify important changes that could drive the decreased innate immune response(s) and potentially identify novel biomarkers for RV16 infected epithelial cells. We challenged HeLa-Ohio cells with RV16 at an MOI 1 for either 6 or 18 hours and at each time point collected the supernatants and processed them for miRNA analysis using a custom panel of 88 different miRNAs. Following qPCR, the results were imported into the GeneGlobe software for further analysis. Our results found that out of the 88 different miRNAs, 9 were modified across our time course with differences in the pattern of regulation (Figure 4). First, we found that 3 miRNAs were upregulated at 6 hours and then downregulated by 18 hours (hsa-let-7f-5p, hsa-miR-101-3p, hsa-miR-34a-5p and hsa-miR-374a-5p) (Figure 4A, Table 3). Secondly, we found that 4 different miRNAs were upregulated at 6 hours and further upregulated by 18 hours (hsa-let-7i-5p, hsa-miR-17-5p, hsa-miR-181b-5p and hsa-miR-30b-5p) (Figure 4B, Table 3). Finally, we found that 2 miRNAs were either upregulated at 6 hours and then downregulated (hsa-miR-21-5p) or only upregulated at 18 hours (hsa-miR-424-5p) (Figure 4C, Table 3). Taken together these results suggest firstly that RV16 infection of HeLa-Ohio cells modifies the miRNA landscape, and this is dependent on time point and productive infection. Secondly, our results suggest that a panel of these miRNA could be useful biomarkers for confirmation of RV16 infection and possible estimation of the infection time.

**Table 3.** Identified miRNAs and their fold changes.

| miRNA | Fold change at 6h | Fold change at 18h |
| --- | --- | --- |
| *hsa-let-7i-5p* | 6 | 16 |
| *hsa-miR-101-3p* | 19 | 4 |
| *hsa-miR-17-5p* | 5 | 9 |
| *hsa-miR-181b-5p* | 3 | 6 |
| *hsa-miR-30b-5p* | 3 | 7 |
| *hsa-miR-34a-5p* | 8 | 1 |
| *hsa-miR-374a-5p* | 8 | 1 |
| *hsa-miR-21-5p* | 10 | 3 |
| *hsa-miR-424-5p* | 3 | 12 |

**Figure 4:**
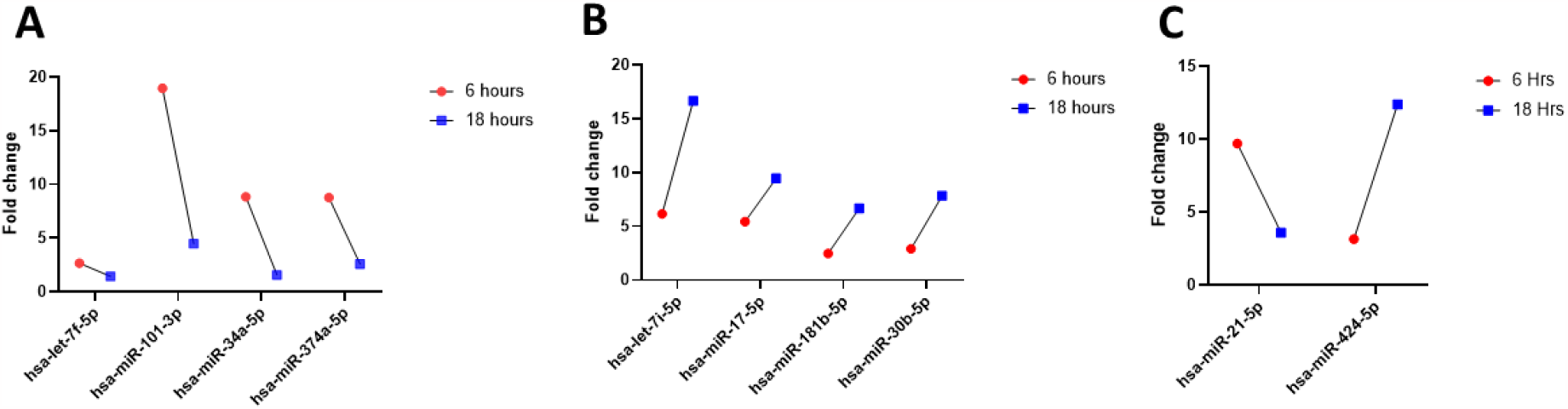
RV16 modifies the expression of miRNAs in infected HeLa-Ohio cells. HeLa-Ohio cells were challenged with RV16, or mock medium at an MOI 1 for 6 or 18 hours and then miRNA expression assessed by qPCR. (A) miRNA with high expression at 6h, (B) miRNA with increasing expression over time, (C) miRNA with increased expression at 6 or 18 h. n=3

### Genes regulated by these microRNAs

To better understand what the miRNA changes meant we next analysed pathways controlled by those miRNAs using the Kyoto database. From these results we were able to create a heatmap (Figure 5) that showed a plethora of the identified miRNAs were proven to play a role in regulating immune pathways responding to viral infection (Figure 5). Specifically, we found that there was significant involvement of hsa-miR-34a-5p in multiple pathways including viral carcinogenesis (-log (Value)>10) specifically in response to HTLV1 infection (-log(P-value)>7.1) and IAV (-log(P-value)>4.8). Additionally, our results showed that other miRNAs e.g., hsa-miR-29a-3p were involved to a lesser extent (Figure 5). These are two examples, but additional miRNAs were also found to play a role including hsa-miR-101-3p. This suggests that the miRNAs we highlighted in figure 4 as well as others do play a role in the innate immune response to viruses and therefore could control genes responsible for our phenotype with RV16.

**Figure 5:**
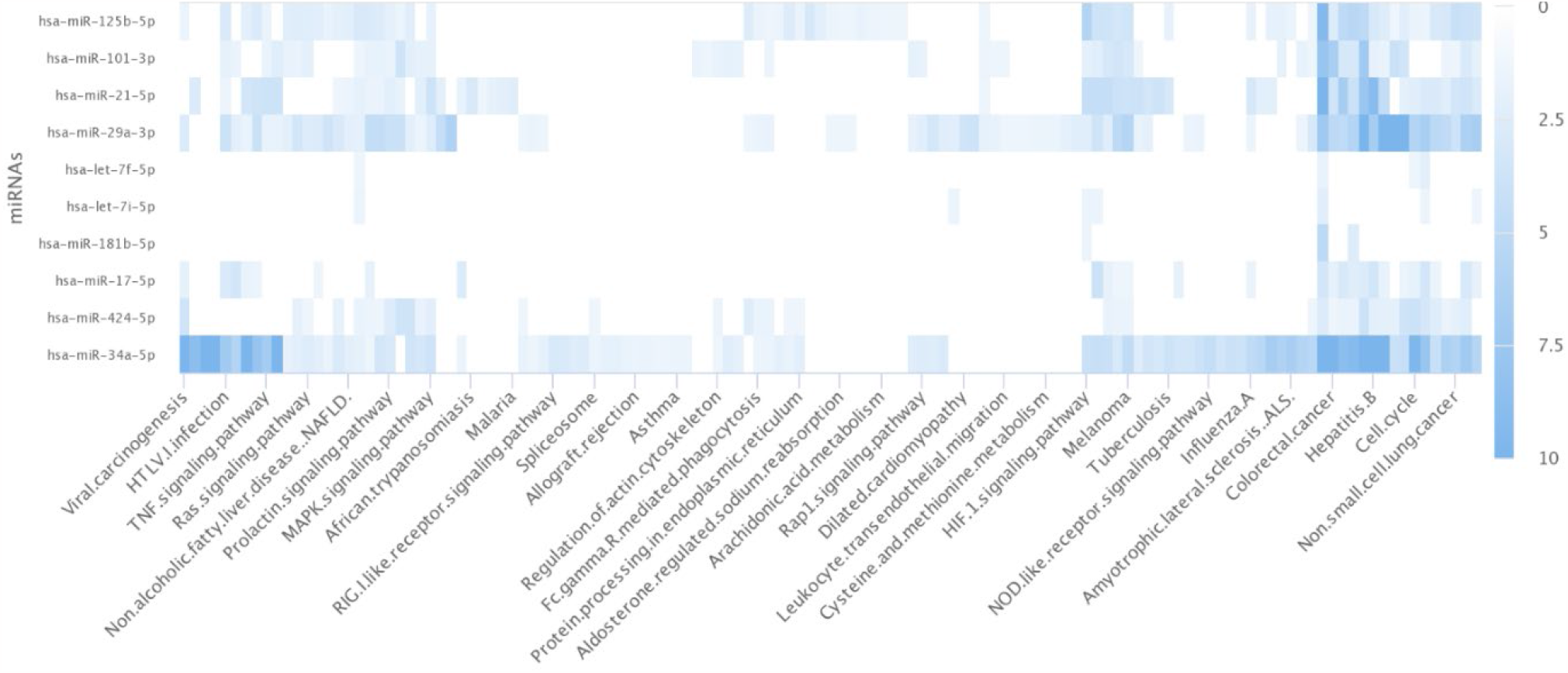
A subset of identified miRNAs control viral innate immune pathways. The identified miRNAs were imported into the Kyoto Encyclopedia of Genes and Genomes (KEGG) database to identify regulated pathways. This figure represents a heatmap demonstrating the involvement of the identified miRNAs in different innate immune pathways related to viral infection.

### hsa-miR-101-3p and hsa-miR-30b-5p control key genes in the response to RV16

Based on the above we next developed a computer program to analyse which miRNAs were controlling a specific phenotype-the response to RV infection and related downstream genes. The programme was developed as outlined in the materials and methods. The identified miRNAs in figure 4 were used as an input variable in the parameters for the API request to mirDIP database. The mirDIP database returned results containing the genes they regulate, and the score of that relationship. Due to the massive amount of returned data, the top three results for each miRNA were chosen for the next step. The chosen genes became a part of the parameters for the VarElect API request. The VarElect database returned results containing the genes that were identified as having a direct connection to our chosen phenotype-innate immune response to RV infection (Figure 6A). From this analysis, first using mirDIP we identified a range of different genes that were related to the different miRNAs we identified in figure 4 (Figure 6A). Taking this further using VarElect we were then able to determine specific genes related to our phenotype controlled by these miRNAs. From this VarElect analysis we were able to identify a panel of different genes and selected the three genes that scored the highest and corresponded to our phenotype (EZH2, RARG and PTPN13) (Figure 6B) and found that these were controlled by 2 miRNAs (hsa-miR-101-3p and hsa-miR-30b-5p) (Figure 6B). Further analysis found that hsa-miR-101-3p controlled the expression of Enhancer of Zeste 2 Polycomb Repressive Complex 2 Subunit (EZH2) and is downregulated when the miRNA is upregulated (Figure 6B). Secondly, we found that hsa-miR-30b-5p specifically downregulated RARG and PTPN13 at later time points (Figure 6B). Together these results suggest that both miRNAs and their small number of RV specific downstream genes could potentially serve as biomarkers for RV16 infection of epithelial cells.

**Figure 6:**
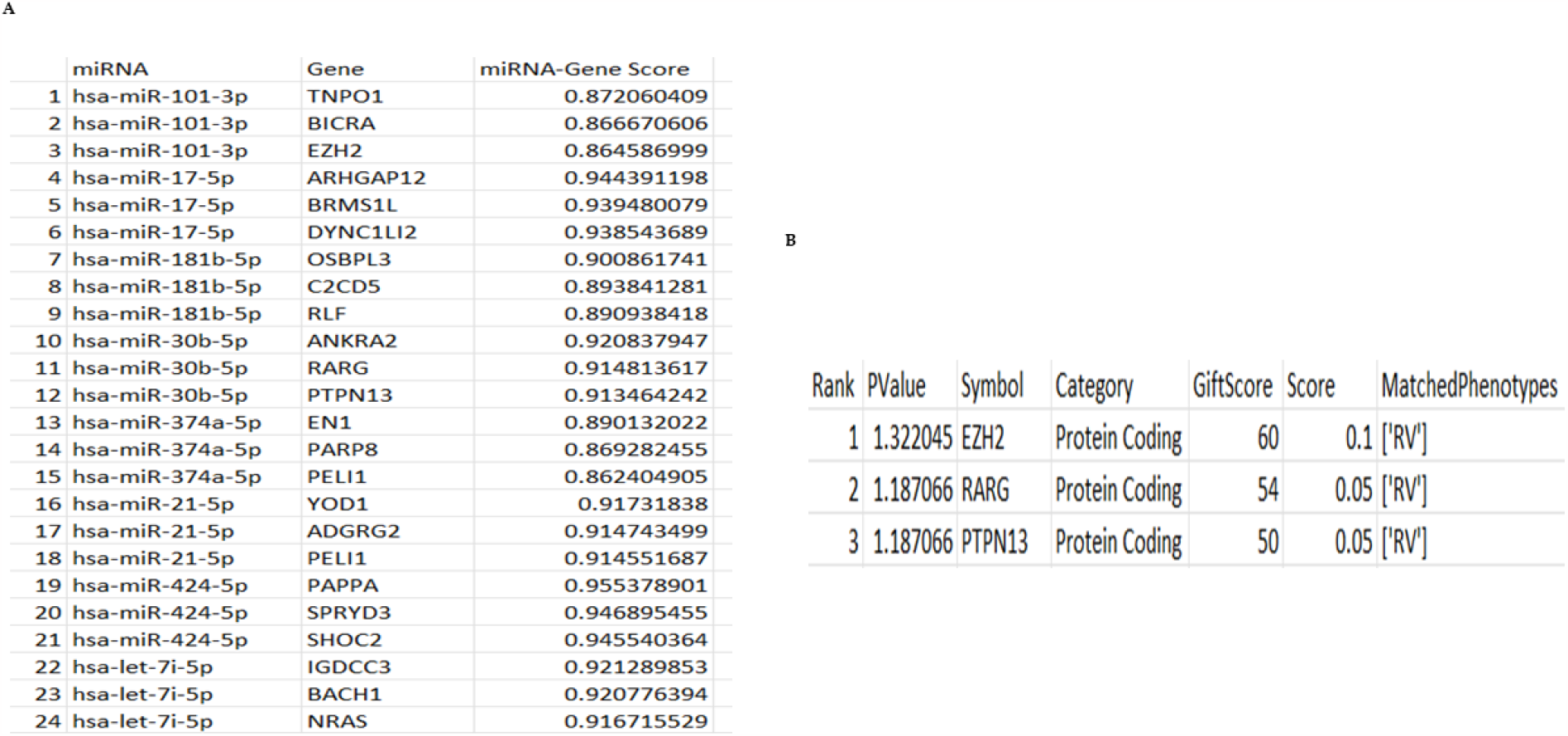
*hsa-miR-101-3p* and *hsa-miR-30b-5p* regulated genes control the response to RV. The identified miRNAs were imported into our newly developed programme to identify which miRNAs were regulating genes responsible for our RV phenotype. Results were all generated using mirDIP and VarElect (A) Table showing all miRNAs and regulated genes based on the formatted mirDIP results and (B) Three most signficant genes identified from the analysis using the formatted VarElect results.

## Discussion

In this study, we show that RV16 prevents complete *S. aureus* clearance over a short timeframe by HeLa-Ohio cells allowing the bacteria to replicate intracellularly and identify potential genes that could be modified by RV16 to impact the host immune response driving this phenotype. We identify that RV16 modifies the miRNA landscape in HeLa-Ohio cells differentially over time. We further identify for the first time using a newly developed program which miRNAs are the most likely candidates driving our phenotype and have identified three likely candidate genes that could promote RV16 infection within HeLa-Ohio cells and potentially compromise the overall immune response. Overall, our results have produced a new program that can be used to analyze miRNA and downstream correlated genes in different scenarios and has identified two potential miRNA biomarkers of RV16 infection. The identification of these two different miRNAs, modified at different time points of RV16 infection represents a potentially significant finding for the field. Our study has potential implications as it suggests that miRNAs modified by RV16 could be used as potential screening and diagnostic biomarkers for at risk patients which could lead to new and better treatments being discovered which could eventually decrease disease burden and improve the overall quality of life.

Our understanding of how RV infections alter epithelial cells leading to disease exacerbations remains inconclusive. Furthermore, there is currently no universal treatment for RV infections [21] and no clear predictive biomarker of infection despite some early excitement around IP-10 [22]. In our work we first set out to determine whether RV16 alters the ability of epithelial cells to respond to bacterial targets in a similar manner to macrophages [8, 21]. To our surprise, despite a robust and productive infectious cycle, HeLa-Ohio cells were not compromised in their ability to control the internalization of two different bacteria that can drive disease exacerbations [6]. It is possible that this normal level of internalization could represent increased cell death in the infected population, but our initial findings would not suggest this. Furthermore, it could suggest increased binding of bacteria to the HeLa-Ohio cells which can occur under conditions of stress [23] but our prior work ruled this out. Taken together this emphasizes that RV16 does not alter the internalization step of phagocytosis in HeLa-Ohio cells. This is important in the context of diseases such as asthma and chronic obstructive pulmonary disease (COPD) where RV16 is frequently isolated [6] and suggests that a key cell type involved in the initial response to infection in the airways is still able to respond in the presence of an ongoing RV16 infection. However, normal levels of internalization do not always translate into a complete immune response (Jubrail et al 2023 in revision) and downstream defects can also drive later bacterial infections [24-25].

Downstream of bacterial internalization, a coordinated series of steps occurs eventually culminating in phagolysosome formation [26-28]. Within this phagolysosome bacterial clearance occurs through various mechanisms [29] and numerous bacteria have been shown to evade these mechanisms [25, 30]. Therefore, we next determined whether following RV16 infection, HeLa-Ohio cells were able to clear intracellular *S. aureus*. Interestingly, despite the initial rapid phase of bacterial killing in infected cells which reduced bacterial numbers to the same as uninfected cultures (over 2 hours), this failed to be sustained over time. This initial phase of rapid killing post phagocytosis has been documented in macrophages and *in vivo* [25] and is usually suggestive of NADPH oxidase activation [31]. When we assessed the percentage killing in our cultures, we showed this to be greater in infected cells in the first 30 minutes which then slowed over the next 2 hours compared to control cultures that almost eradicated the intracellular inoculum. To our surprise, in the RV16 infected cultures, beyond 2 hours, intracellular bacterial numbers began to increase. This is suggestive of intracellular bacterial replication which has been seen in different cell types for *S. aureus* [24-25, 32]. Despite these earlier, reports we are the first to observe this phenomenon in the context of secondary bacterial trigger post-viral infection and the first to observe this at such at early time point, with earlier studies suggesting at least 3 days is required before the bacteria begin replicating. It is possible that this is because the RV16 infection has sufficiently altered intracellular organelles in infected cells which supports bacterial replication in immature phagolysosomes [25] or that phagolysosome maturation is stalled allowing the bacteria to replicate in the cytoplasm which has been observed for *S. aureus* in macrophages [33]. In the context of diseases such as asthma and COPD this is an important finding because increased staphylococcal replication could lead to an overt epithelial cell death perpetuating inflammation within the airways. The released bacteria could then be reinternalized by macrophages which are known to act as Trojan horses of *S. aureus* infection predisposing individuals to systemic and life-threatening infections [25, 34].

Based on these findings we decided that we next would begin to decipher if RV16 infection modified the global miRNA landscape in HeLa-Ohio cells with a view to both uncovering novel miRNAs that linked to our prior findings and to identify potential biomarkers of RV16 infection. In our studies we used a custom panel of 88 miRNAs and assessed their expression at 6- and 18-hours post RV16 infection. From this we identified a panel of 9 miRNAs demonstrating differential expression over our time course with a panel being initially upregulated and then downregulated by 18 hours, another panel continually increasing in expression over time and a final panel being up and then downregulated or vice versa. This pattern of miRNA expression could be representative of a dynamic interaction between the virus and host with both positive and negative impacts for viral replication and a protective immune response as has been shown in other infections [35-37] and in our initial analysis using the KEGG database. Previous studies assessing the impact of RV on miRNA expression have identified limited miRNA’s that could be important for ongoing infection such as miR-155 and miR-128 [15-16]. Unlike these previous studies, our work has used two timepoints of infection and demonstrated how the miRNA landscape changes over this time course rather than remaining static. In the context of airway infection and inflammation our work represents an important advance because it has highlighted a subset of miRNAs that could represent screening biomarkers in at risk patients and a second subset of miRNAs that could be future diagnostic biomarkers for patients with diseases such as asthma and COPD who suffer with ongoing RV exacerbations [6]. Overall, these results showed that RV16 does modify the miRNA landscape in infected HeLa-Ohio cells and that this could correlate with the timing of infection.

To better understand the impact of these changes on HeLa-Ohio cells we next set out to understand if any of our identified miRNAs could control our previous findings with *S. aureus* and represent true biomarkers for RV16 infection. To do this we developed a new computer modelling program that would allow us to both correlate the genes associated with a panel of miRNAs and then to specifically identify genes associated with a phenotype we were interested in. Our initial analysis of this demonstrated that all the identified miRNAs in our panel were responsible for controlling the HeLa-Ohio response to RV16 infection. However, when we then analyzed the data further, we were able to identify three specific genes that were the strongest represented within our dataset correlated to two specific miRNAs, miR-101-3p and miR-30b-5p. These miRNAs represent two different subsets within our data with miR-101-3p being increased ∼20 fold at 6 hours and then decreasing over time and miR-30b-5p showing relatively modest induction at 6 hours and increasing to ∼8 fold at 18 hours. Based on this analysis we strongly believe that miR-101-3p could represent a good screening biomarker for RV16 infection and miR-30b-5p could represent a good diagnostic biomarker for RV16 infection.

This idea of miRNA’s being biomarkers has gained traction in recent years [10] and has recently gained further popularity with IAV infection [37]. In our work we found that increased expression of either of these two miRNAs correlated with a possible promotion of viral infection (Figure 5) at the expense of the immune response. Importantly, our program linked to VarElect identified that miR-101-3p controlled the expression of EZH2. Although this has never been explored in RV infections it has been documented to drive a potent antiviral state and maintain immune homeostasis in cells when it is downregulated and miR-101-3p controls its expression levels [38]. This supports our data and suggests that at early time points post infection when viral replication is low host cells could sufficiently induce an antiviral state through miR-101-3p and EZH2. Over time however, as viral replication increases and the expression of miR-101-3p decreases this could correlate with a promotion of viral replication and spread and a weakened host antimicrobial response supporting our earlier findings. Importantly a recent study showed in T cells that RV can manipulate the expression of EZH2 driving T cell apoptosis and enhanced viral replication [39]. However, this study failed to correlate this with miRNA expression and furthermore reported viral replication in T cells which has always been a matter of debate. Our analysis further identified that miR-30b-5p controlled two further genes important in the immune response RARG and PTPN13 whose expression decreased when the miRNA expression increased. There have been few studies correlating these genes with viral infections, but the limited literature does associate them with more severe outcomes particularly PTPN13 [40]. In the case of RARG, it’s been suggested it could play key roles in the response to *S. aureus* meaning any downregulation could potentially have a detrimental effect on the immune response [41]. Clearly being able to understand more about these genes and their role in viral infection will be beneficial. Overall, these results do suggest two key miRNAs and their downstream genes could function as novel biomarkers of RV16 infection.

Our study whilst a step forward does have a few limitations. First, we performed our work in HeLa-Ohio cells and therefore need to extrapolate these findings to primary epithelial cells in the future. Secondly, we used a major group RV and therefore seeing if these findings also hold true with a wider panel would be interesting. Thirdly, we haven’t in this study verified the downstream genes controlled by each miRNA and this remains the focus of future papers. However, we aimed to identify a predictive panel of miRNAs and show that our new program can be used to quickly and reliably identify the genes associated with these miRNAs, which represents a major methodological advance.

Overall, given the limited information on the role of miRNAs during RV infection, our work represents a step forwards. Not only have we identified changes in the miRNA landscape that could link viral infection to downstream responses, but we have identified a panel of miRNAs that could be potential diagnostic and screening biomarkers. Within this we suggest that miR-101-3p could represent a good screening biomarker and miR-30b-5p could represent a good diagnostic biomarker. These biomarkers can aid in early diagnosis of RV infections, preventing patients suffering long term exacerbations. Furthermore, as miRNAs are unique, it could be possible to incorporate them and their response elements into a novel RV vaccine along with key immunogenic viral proteins. Additionally, our future work will examine the effect of targeting these miRNAs which could lead to the generation of new and specific RV therapeutics. In the future, we will need to explore the effect of these miRNAs and the downstream genes on the viral life cycle and secondary infections with more mechanistic experiments to better understand their overall impact and importance during RV infection.

Finally, these results show that some miRNAs play an important role during RV16 infection particularly when the virus is actively replicating and assembling in epithelial cells. The two miRNAs we identified in this study may be of diagnostic value as their expression levels differ during different stages of infection. This may lead in the future to developing a new diagnostic test and potential treatment for RV16 infection that can be used globally but will be of particular importance for patients where this virus causes high mortality.

## Conclusions

In conclusion, the present study examined the impact of RV16 on both epithelial cell antimicrobial responses and the miRNA landscape. The results identified that RV16 infection of HeLa-Ohio cells promoted staphylococcal replication and that a panel of miRNAs correlated with immune defenses which could explain these results. The results provide the potential that two miRNAs, miR-101-3p and miR-30b-5p could be potential biomarkers for RV16 infection and could help explain our earlier findings. These results expand on the current miRNA landscape work in RV infections and further show how the virus can control host gene expression. Our current results descriptively suggest that miR-101-3p and miR-30b-5p could be two different types of biomarkers for RV16 infection and further emphasize the need to understand with more mechanistic detail the role of miRNAs and their related genes during RV infection of multiple cell types.

## Author Contributions

“Conceptualization, JJ. and AK.; methodology, PB, VC, AT, AK and JJ.; software, PB.; validation, PB, AK and JJ.; formal analysis, PB, VC and AT.; investigation, PB, VC and AT.; resources, AK and JJ.; data curation, PB, VC and AT.; writing—original draft preparation, PB, VC, AT and JJ.; writing—review and editing, all authors.; visualization, all authors.; supervision, AK and JJ.; project administration, AK and JJ.; funding acquisition, AK and JJ. All authors have read and agreed to the published version of the manuscript.”

## Data Availability Statement

All data generated in this manuscript will be made available upon request after consulting the database providers if needed.

## Acknowledgments

The team would like to thank Dr. Amr Abdelgany and Dr. Hannah Rickman for technical expertise, essential laboratory training for PB, VC and AT and for insightful discussions about the data. Thanks, are also extended to the Weizmann Institute of Science for granting us the right to use the VarElect tool (GeneCards database) which is an integral part of the developed program.

